# *tidygenclust*: Clustering for Population Genetics in R

**DOI:** 10.1101/2025.07.29.667403

**Authors:** Eirlys E. Tysall, Anahit Hovhannisyan, Evelyn J. Carter, Cecilia Padilla-Iglesias, Margherita Colucci, Andrea Vittorio Pozzi, Michela Leonardi, Aramish Fatima, Ondrej Pelanek, Nile P. Stephenson, Andrea Manica

**Affiliations:** Department of Zoology, University of Cambridge, Cambridge, CB2 3EJ, United Kingdom; Smurfit Institute of Genetics, Trinity College Dublin, Dublin, D02 PN40, Ireland; School of Life Sciences, Anglia Ruskin University, Cambridge, CB1 1PT, United Kingdom; Emmanuel College, University of Cambridge, Cambridge, CB2 3EJ, United Kingdom; Human Palaeosystems Group, Max-Planck Institute of Geonanthropology, 07745 Jena, Germany; Natural History Museum, London, SW7 5BD, United Kingdom; University Museum of Zoology, University of Cambridge, Cambridge, CB2 1RB, United Kingdom

**Keywords:** clustering, tidypopgen, fastmixture, Clumppling, admixture, R, population genetics, population genomics, population structure, bioinformatics

## Abstract

**Background:** Population structure analysis is crucial for evolutionary research and medical genomics. Clustering methods, broadly categorized as model-based (e.g. ADMIXTURE) or non-model-based (e.g. SCOPE), differ in their methodology and computational efficiency. Recently, *fastmixture*, a model-based approach, has improved scalability and performance, while replicate alignment tools, such as Clumppling, extend previous methods by also aligning the modes across K values. However, all the existing tools are standalone and generate numerous untracked text files, as well as offering limited plot customisability.

**Results:** We introduce an R package, *tidygenclust*, which brings the functionalities of the original ADMIXTURE, *fastmixture* and Clumppling software into R, enabling a streamlined and integrated workflow. By integrating with *tidypopgen*, a package designed to handle large SNP datasets, these new tools maintain metadata, simplify data handling, and produce results as customisable *ggplot2* objects for flexible visualisation.

**Conclusions:** The R package *tidygenclust* advances population genetic analysis by combining computational efficiency with reproducible workflows and user-friendly plotting. The source code and instructions can be accessed on https://github.com/EvolEcolGroup/tidygenclust.

## Background

Understanding population structure is essential for evolutionary studies and medical genomics. Clustering methods to infer population structure can be broadly divided into two main categories: model-based and non-model-based approaches. Model-based methods, which include the widely used and considered the gold standard ADMIXTURE [1], rely on probabilistic models to estimate ancestry proportions but often struggle with scalability when applied to large datasets. In contrast, non-model-based methods use generic statistical approaches, such as principal component analysis [2], and provide a more computationally efficient alternative, especially for large datasets, but they may not offer the same level of detailed insights into population structure. A recently developed model-based method, *fastmixture* [3] combines improved scalability and accuracy for modern genomic studies.

A common challenge with unsupervised clustering methods is the variability in output memberships when clustering the same set of individuals across different runs due to stochasticity. To address this issue, several methods have been implemented, employing various algorithmic approaches for the alignment of replicates, such as Clumpp [4], Clumpak [5], and Pong [6]. An advancement in replicate alignment has been introduced with Clumppling [7], which extends previous methods by also aligning the modes across K values and, importantly, improves computational time.

However, both clustering and replicate alignment software have their limitations when it comes to providing user-friendly outputs and tracking the provenance of data. While other population genetics analyses have been integrated into pipelines, genetic clustering has generally been in the form of standalone software. This applies to standard clustering methods, such as ADMIXTURE, which is freely available but not open source, and more recent developments such as SCOPE, a C++ command line program, and *fastmixture*, a Python-based software compiled as a command line program to be used in bash. Users must use custom scripts to extract relevant data from the multiple files generated by these tools. Similarly, replicate alignment software tools are self-contained and generate numerous text files with no metadata for tracking. Additionally, these programs create a predetermined set of plots, rather than outputting a plotting object, which can be modified by the user. A notable exception is openADMIXTURE [8], which is coded in Julia and interfaces with libraries in that language. However, since Julia is not widely used in population genetics, this poses a challenge for its adoption. These constraints, combined with the lack of integration between clustering and replicate alignment programs, often prevent users from performing multiple clustering runs.

Genetic clustering has received relatively little attention in R, except for a number of functions in various packages to plot the results of externally run software. A notable exception is sNMF in the package LEA [9], which provides clustering based on sparse non-negative matrix factorisation (an earlier approach that bears several similarities with SCOPE). However, this is simply a port of the original standalone C program so that it is compiled directly in R, and it still requires creating a custom input text file for the genotypes. Snapclust [10] provides a fast likelihood clustering method fully integrated into the commonly used R package *adegenet*, but the Expectation-Maximisation algorithm does not scale to large datasets. The features of both genetic clustering and alignment software are summarised in Table 1.

**Table 1.**
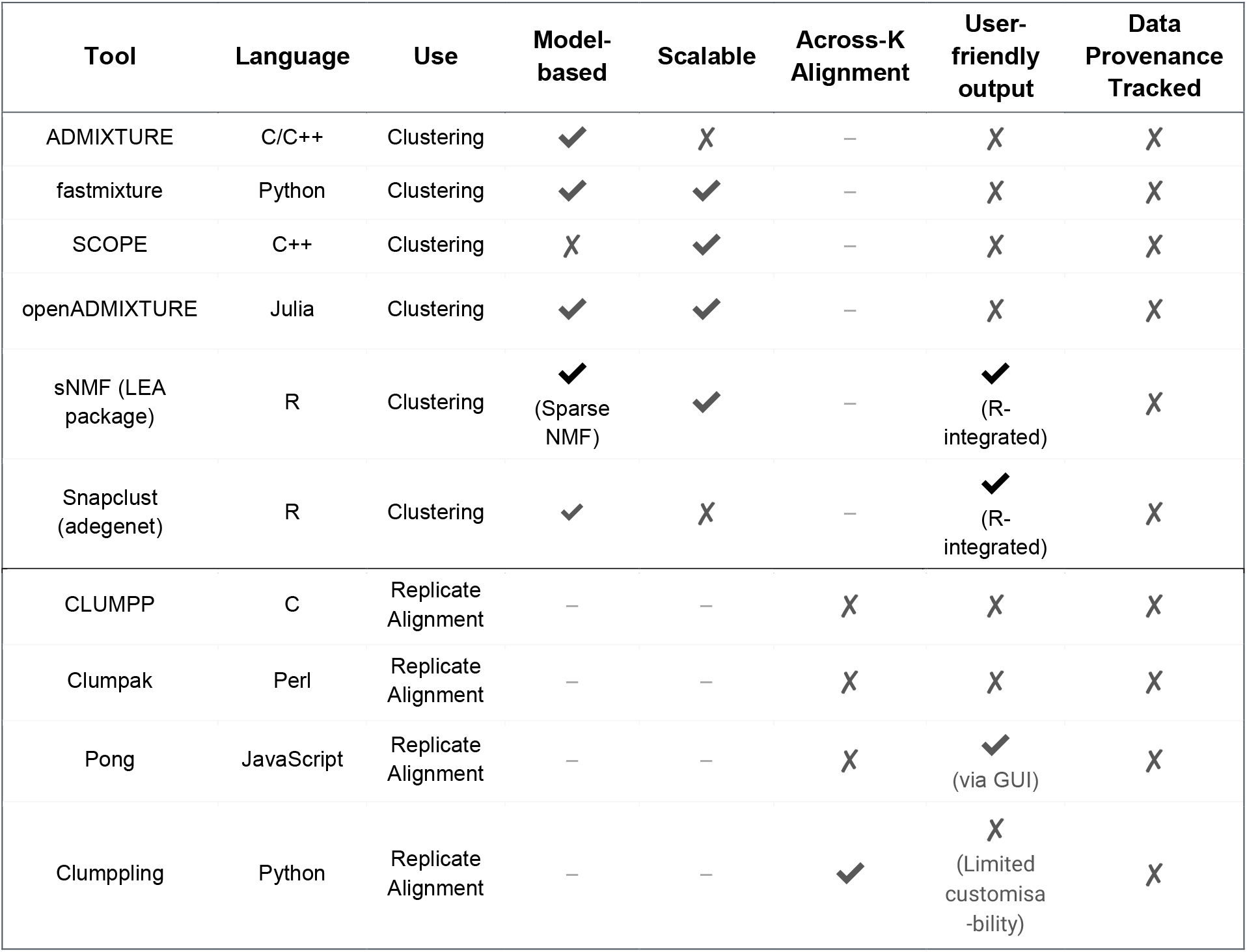
Summary of the features of currently available software for clustering analyses and replicate alignment.

## Implementation

### Implementation

Recently, we developed the *tidypopgen* package [11] for the manipulation and analysis of genetic data in R. It allows users to work with native objects and easily transition from one analysis to another while carrying their metadata. Here, we present an additional package for population genetic clustering directly in R—*tidygenclust*, which can be fully integrated into *tidypopgen* pipelines. *tidygenclust* provides R interfaces to the original Python-based and standalone pieces of software *fastmixture* and Clumppling. We used the reticulate package [12], to enable seamless passing of objects between R and Python. While the functions can work with text files, as the original software, they also provide interfaces to work directly with R objects from *tidypopgen*. This enables users to run all analyses entirely within R, with clear tracking of data provenance stored in coherent data structures, avoiding the creation of numerous text files that are difficult to manage and lack full metadata. Additionally, this approach allows for the easy editing and customisation of plots.

### Details of the supported standalone software

**ADMIXTURE** [1] is a model-based software for population genetic clustering. It performs a maximum likelihood estimation for individual ancestries on SNP genotype datasets. It is arguably the most widely used tool for genetic clustering, due to its reliable performance and reasonable speed.

***fastmixture*** [3] is another model-based software for population genetic clustering. It uses randomized singular value decomposition and a mini-batch accelerated scheme to improve the convergence of the expectation-maximization algorithm, significantly enhancing speed in ancestry estimation compared to the ADMIXTURE software. Compared with non-model clustering approaches, *fastmixture* enables more accurate ancestry proportion estimation for larger sample sizes and whole-genome sequencing data, making it a promising tool for population genomic and genome-wide association studies.

**Clumppling** [7] is a method for aligning replicate unsupervised clustering runs, utilizing integer linear programming for optimal alignments within combinatorial optimization frameworks. It builds on previous tools, extending their capabilities by enabling alignment across multiple K. Compared to the previous methods, Clumppling delivers higher similarity scores and faster computation times, making it a more efficient and effective approach for aligning clustering results.

### Source code and software availability

*tidygenclust* is written in R, and the source code is available free online on https://github.com/EvolEcolGroup/tidygenclust. It can also be installed directly from the R-Universe https://evolecolgroup.r-universe.dev/tidygenclust. The package works on Linux and OSX, and can be used on Windows via the Windows Subsystem for Linux (WSL2).

## Results

Here, we demonstrate how to analyse data using the *tidypopgen* and *tidygenclust* packages in R. As an example, we use ~1000 individuals from 52 populations from the HGDP dataset [13]. The dataset is first imported into R as a *gen_tibble* object. We then perform some basic quality filtering for missingness and monomorphic loci before running linkage disequilibrium (LD) pruning using functions within *tidypopgen*. On this filtered and LD pruned ‘gen_tibble’ we run ‘gt_fastmixture()’ for one repeat run of K = 5. The result is provided in the form of a ‘gt_admix’ object which stores all the clustering outputs; Q matrices, P matrices, log files and cross-validation where possible (this is available for e.g. ADMIXTURE), for each run of K alongside any population or grouping meta-data provided in the ‘gen_tibble’. We can autoplot the result from the ‘gt_admix’ object directly using *tidypopgen* (Figure 1).

**Figure 1.**
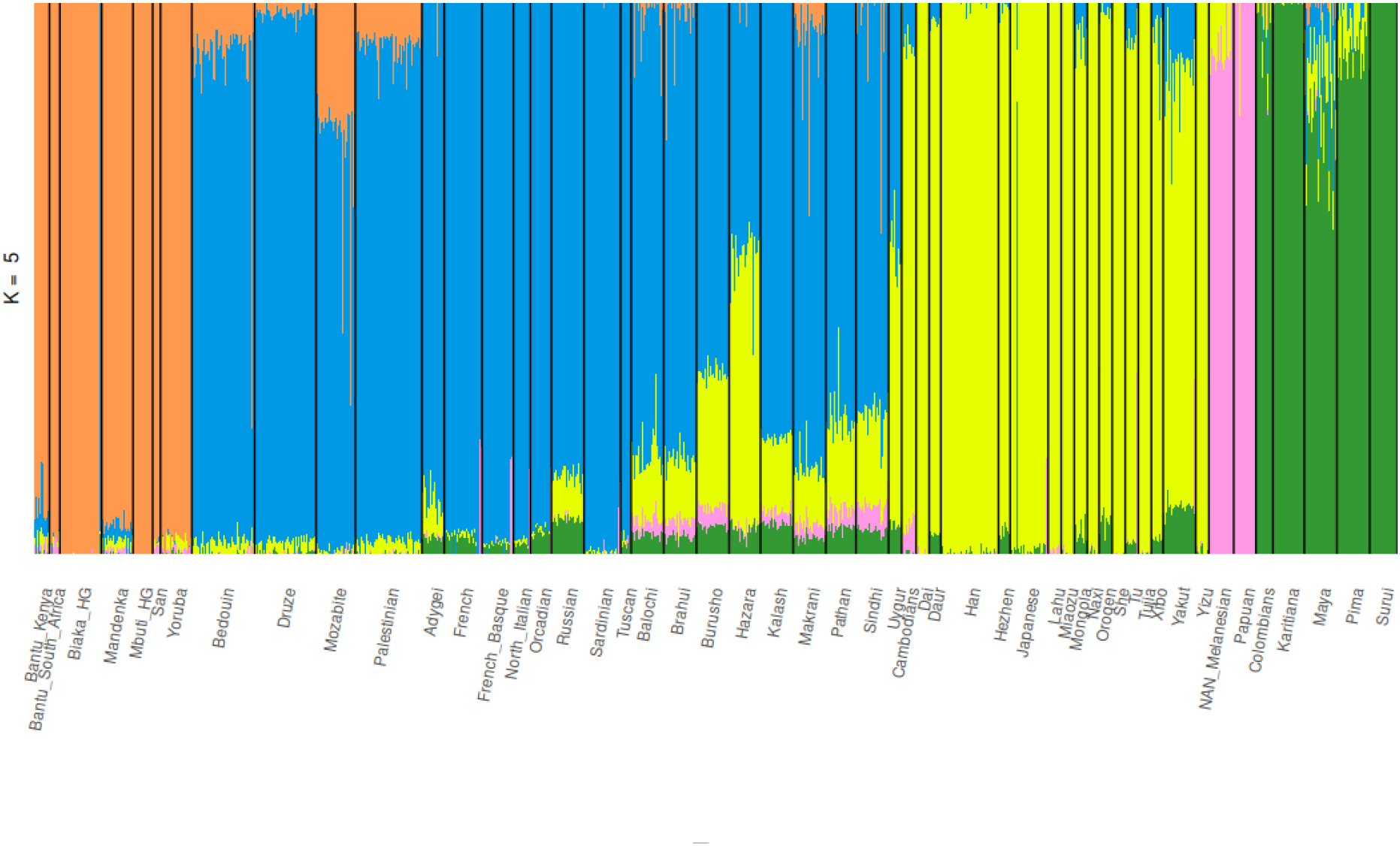
Autoplot of ancestry proportions identified using the ‘gt_fastmixture()’ function to run *fastmixture* in *tidygenclust* for one run of K = 5 on the HGDP dataset.

**Figure 2.**
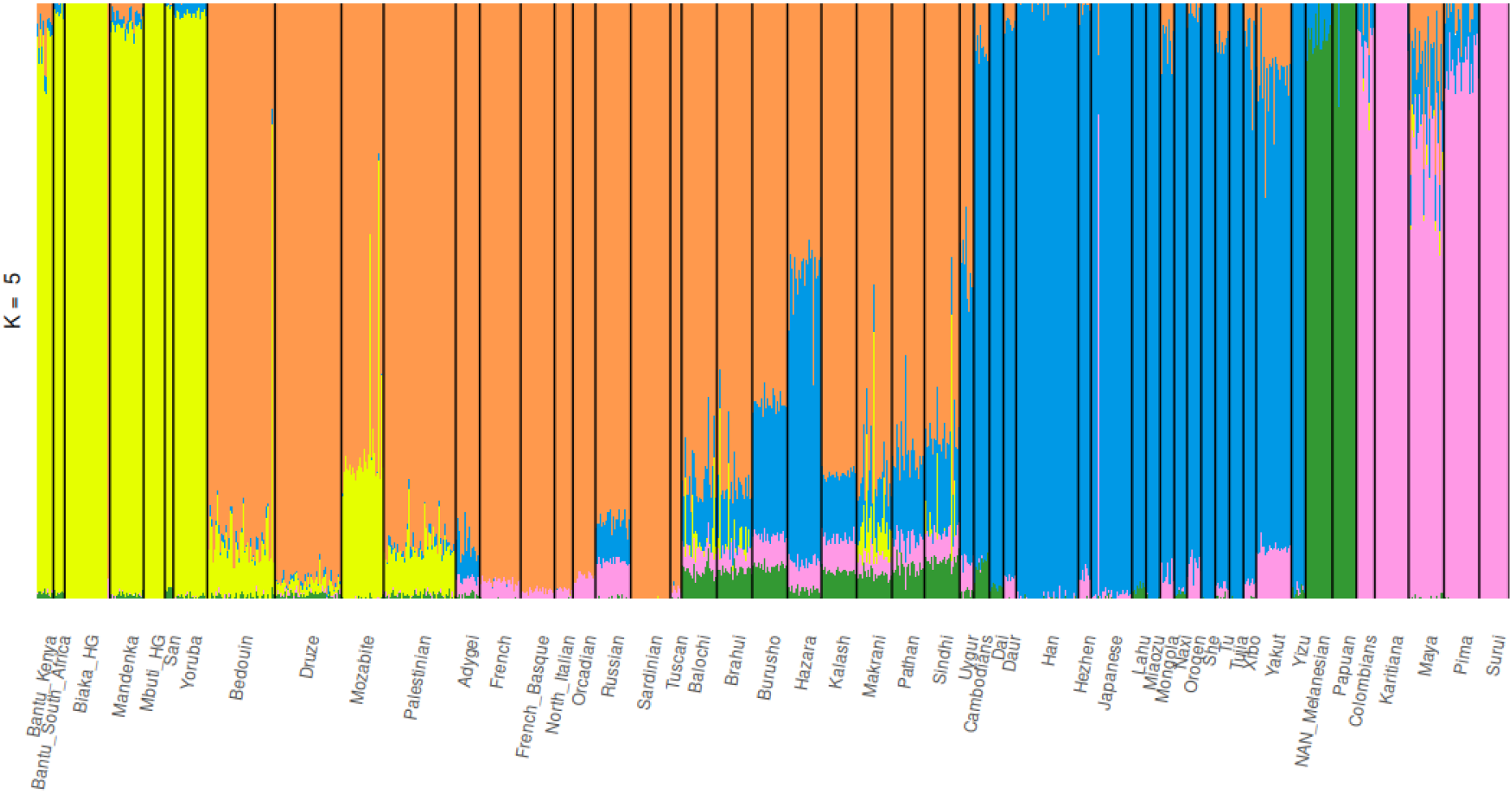
Autoplot of ancestry proportions identified with ADMIXTURE within *tidypopgen* for one run of K = 5 on the HGDP dataset.

We compare the results of *fastmixture* within *tidygenclust* to those obtained from running ADMIXTURE within a convenient wrapper, ‘gt_admixture()’ which is part of *tidypopgen*. Whilst ADMIXTURE usually requires PLINK files as input, here ADMIXTURE is directly integrated into the *tidypopgen* framework and so we can use the same ‘gen_tibble’ object as input, and the output is another ‘gt_admix’ object with the same structure as the one obtained from ‘gt_fastmixture()’.

Having compared outputs between *fastmixture* and ADMIXTURE, we extend our analysis to a broader range of K values. Firstly, we use ADMIXTURE’s cross-validation procedure to assess what might be the most appropriate value of K for our dataset. We run ADMIXTURE in cross-validation mode for K = 2 to 15. It is possible to run all simulations with a single command ‘gt_admixture()’ which returns an object of class ‘gt_admix’ that contains the outputs for all K, including the cross-validation scores. We can plot the latter directly using autoplot (Figure 3).

**Figure 3.**
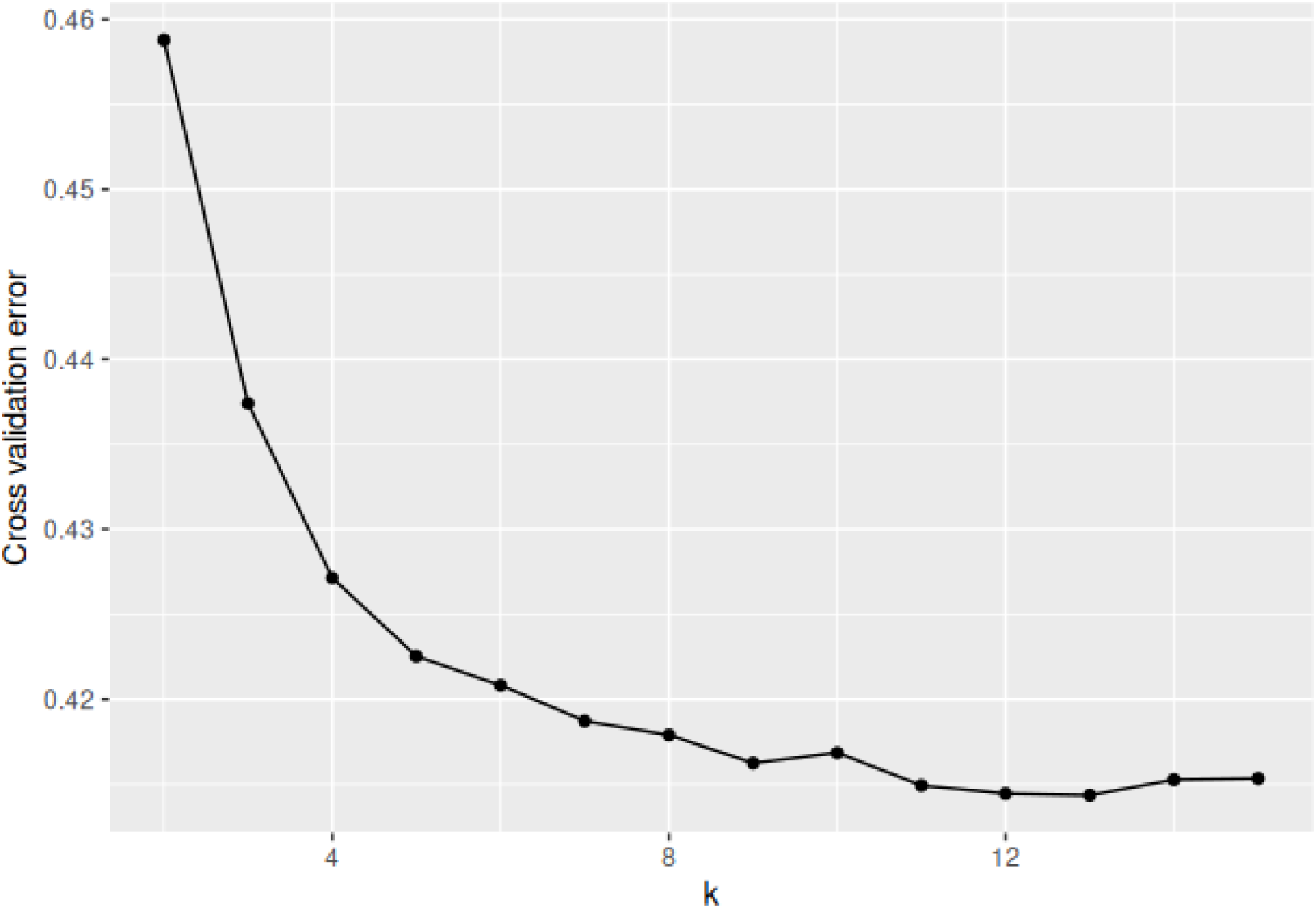
Autoplot of cross-validation scores from running values of K = 2 to 15 in ADMIXTURE within *tidypopgen* on the HGDP dataset.

Next, we perform multiple runs of *fastmixture* using *tidygenclust* for several K values from 4 to 8 with 50 repeats of each value of K. Again, this is all achieved by issuing a single ‘gt_fastmixture()’ command with the appropriate parameters defining the values of K and the number of repeats. We then supply the ‘gt_admix’ results object directly to ‘gt_clumppling()’ within *tidygenclust* to summarise and plot the clustering results. ‘gt_clumppling()’ uses Clumppling to align and summarise the modes of the multiple runs for each K, and the relationship among modes. Firstly, we visualise all the possible modes for each value of K and align these in a single multipartite plot using the ‘autoplot()’ function (Figure 4).

**Figure 4.**
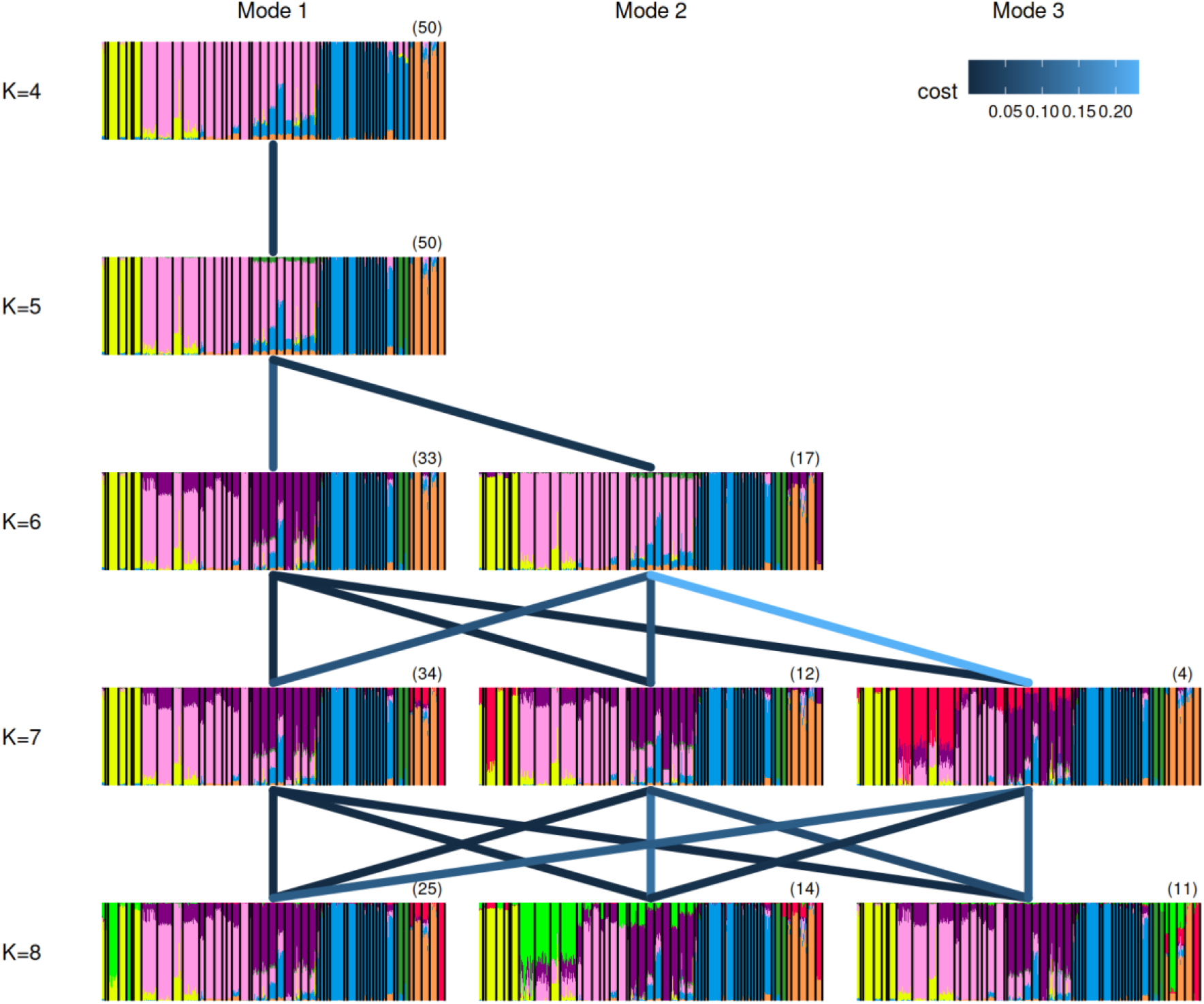
Autoplot of Clumppling aligned modes of 50 repeat runs of K = 4 to 8 as estimated by *fastmixture*. Modes across different values of K are connected by edges, where darker colours indicate larger weights, corresponding to better alignments between modes. The number of runs corresponding to a mode is indicated in brackets above each plot.

Multiple autoplots are available to explore the Clumppling results; we additionally use the “major modes” visualisation which plots only the most supported modes identified by Clumppling for each value of K in a stacked plot (Figure 5). Here we use population groupings carried over from our original ‘gen_tibble’ via the ‘gt_admix’ results object to annotate the Clumppling autoplot plot.

**Figure 5.**
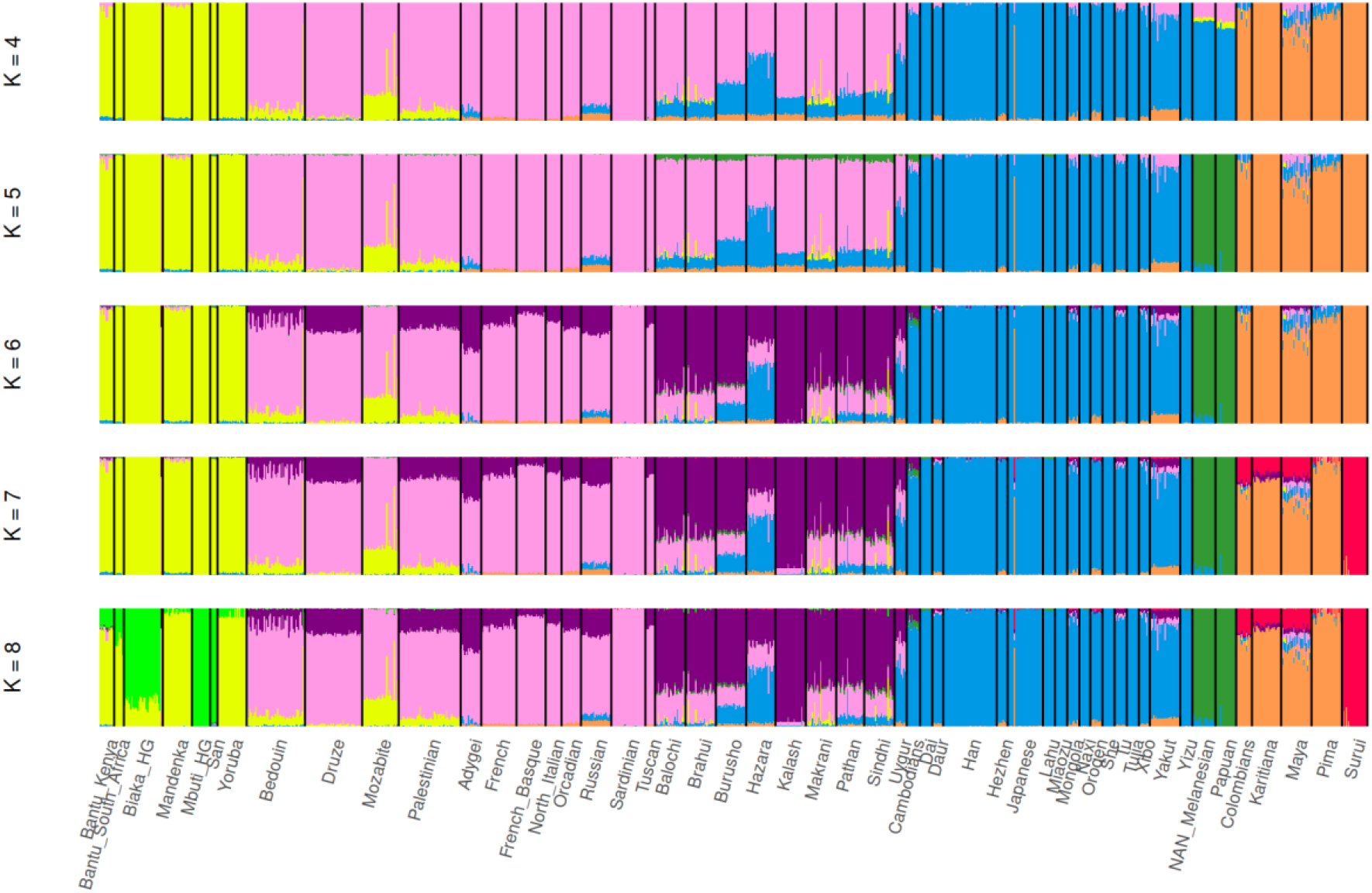
Autoplot of Clumppling aligned major modes of 50 repeat runs of K = 4 to 8 as estimated by *fastmixture*. Only the most supported modes identified by Clumppling for each value of K are shown.

All plots are generated directly in R, as *ggplot2* objects, which can be further edited and modified using the standard R syntax, thus making the preparation of publication quality figures straightforward.

## Conclusions

We present the R package *tidygenclust*, which provides integrated solutions for population genetic clustering and replicate alignment, respectively. This package streamlines the process by enabling users to run all the analyses directly in R, with the added advantage of maintaining metadata, eliminating the need to manage numerous text files in the output as well as having the possibility for editing and customizing the plots.

## Availability and requirements

**Project name:** *tidygenclust*

**Project home page:** https://github.com/EvolEcolGroup/tidygenclust

**Operating system(s):** Linux, OSX, and Windows via the WSL2

**Programming language:** R (and python, via the ‘reticulate’ package in R)

**Other requirements:** None

**License**: GPL3

**Any restrictions to use by non-academics:** None

## Abbreviations

HGDP: Human Genome Diversity Project
SNP: Single Nucleotide Polymorphism
LD: Linkage Disequilibrium

## Declarations

### Ethics approval and consent to participate

Not applicable.

### Consent for publication

The authors consent to the above manuscript being published in BMC Bioinformatics.

### Consent for publication

Not applicable.

### Availability of data and materials

The HGDP dataset analysed during the current study are available in the Zenodo repository, https://doi.org/10.5281/zenodo.15582364. The source code and instructions for *tidygenclust* can be accessed on https://github.com/EvolEcolGroup/tidygenclust and it can be installed directly from R-Universe https://evolecolgroup.r-universe.dev/tidygenclust.

### Authors’ contributions

AM, AH and EET conceptualised and designed the package with input from EJC. AM wrote and implemented the R package with input from EET, AH and EJC. AM and EET wrote the package documentation. AM, EET, AH, EJC, CP-I, MC, AVP, ML, OP, AF and NPS wrote unit tests and improved the package functionality. AH led the writing of the manuscript with input from AM and EET. EJC, CP-I, MC, AVP and ML provided edits and feedback to the manuscript and all authors reviewed and approved the final manuscript.

### Authors’ information

Not applicable.

## Acknowledgements

Not applicable.

